# XMAP215 promotes microtubule catastrophe by disrupting the growing microtubule end

**DOI:** 10.1101/2020.12.29.424748

**Authors:** Veronica Farmer, Göker Arpağ, Sarah Hall, Marija Zanic

## Abstract

The GTP-tubulin cap is widely accepted to protect microtubules against catastrophe. The GTP-cap size is thought to increase with the microtubule growth rate, presumably endowing fast-growing microtubules with enhanced stability. It is unknown what GTP-cap properties permit frequent microtubule catastrophe despite fast growth. Here, we investigate microtubules grown *in vitro* in the presence and absence of the microtubule polymerase XMAP215. Using EB1 as a GTP-cap marker, we find that GTP-cap size increases regardless of whether growth acceleration is achieved by increasing tubulin concentration or by XMAP215. In spite of the increased mean GTP-cap size, microtubules grown with XMAP215 display increased catastrophe frequency, in contrast to microtubules grown with more tubulin, for which catastrophe is abolished. However, microtubules polymerized with XMAP215 have large fluctuations in growth rate and EB1 intensity; display tapered and curled ends; and undergo catastrophe at faster growth rates and with higher EB1 end-localization. Our results underscore the role of growth irregularities in overall microtubule stability.

## INTRODUCTION

Microtubules are cytoskeletal polymers essential for cell motility, cell division, and intracellular transport. A key property of microtubules is that they are highly dynamic, allowing dramatic remodeling of the microtubule network to form cellular structures such as the mitotic spindle. The dynamics of individual microtubules are characterized by ‘dynamic instability’-stochastic switching between phases of growth and shrinkage through transitions known as catastrophe and rescue (Mitchison and Kirschner, 1984). The standard model of dynamic instability implies that the presence of a stabilizing ‘GTP-tubulin cap’ protects a growing microtubule from undergoing a catastrophe. Namely, microtubules polymerize by incorporation of GTP-bound αβ-tubulin heterodimers, followed by the hydrolysis of GTP to GDP in β-tubulin subunits within the microtubule polymer. The balance between the GTP-tubulin dimer addition and subsequent GTP hydrolysis results in a cap consisting of GTP-tubulin dimers at the growing microtubule end. The process of GTP hydrolysis triggers conformational changes in the tubulin dimer, rendering the GDP-tubulin lattice inherently unstable. The loss of the GTP-cap exposes the unstable GDP-tubulin lattice, thus triggering rapid microtubule depolymerization (Mitchison and Kirschner, 1984)(Desai and Mitchison, 1997).

The inability to directly visualize the GTP-cap has made its investigation challenging. Previous studies found that even a small nucleotide cap, consisting of just a few GTP-tubulin layers, can be sufficient to stabilize a growing microtubule end (Drechsel and Kirschner, 1994)(Strothman et al., 2019). Furthermore, an increase in the GTP-cap size, which may occur as a result of an increase in microtubule growth rate, is typically associated with prolonged microtubule lifetime. Along those lines, early work demonstrated that increasing the microtubule growth rate by increasing the tubulin concentration *in vitro* is accompanied by a decrease in the catastrophe frequency (Walker et al., 1988). In recent years, microtubule-associated EB proteins, which display comet-like localization at growing microtubule ends (Bieling et al., 2007), have been established as a marker for the GTP-cap due to their recognition of the nucleotide state of tubulin in the microtubule polymer (Zanic et al., 2009)(Maurer et al., 2012)(Zhang et al., 2015). *In vitro* studies investigating EB localization have revealed that increasing microtubule growth rate by increasing tubulin concentration correlates with larger EB comets (Bieling et al., 2007; Strothman et al., 2019). Additionally, microtubules with brighter EB comets were more stable against catastrophe induced by tubulin dilution (Duellberg et al., 2016). These studies provide further evidence that the suppression of catastrophe at faster growth rates may be a consequence of a larger GTP-cap.

In contrast to microtubules polymerized with purified tubulin *in vitro*, microtubules in cells can simultaneously display fast growth rates and high catastrophe frequency (Rusan et al., 2001)(Mimori-Kiyosue et al., 2005)(Akhmanova and Steinmetz, 2008)(Akhmanova and Steinmetz, 2015). In cells, microtubule dynamics are tightly regulated by a myriad of microtubule-associated proteins (MAPs). Fast microtubule growth rates can be attributed to the action of microtubule polymerases, the most prominent of which are members of the conserved XMAP215 family (Gard and Kirschner, 1987)(Brouhard et al., 2008)(Gard et al., 2004)(Slep, 2009)(Al-Bassam and Chang, 2011). On its own, XMAP215 increases microtubule growth rates up to 10-fold (Vasquez et al., 1994; Brouhard et al., 2008), while a combination of XMAP215 with EB1 synergistically promotes up to a 30-fold increase in microtubule growth rates, matching the fast growth rates observed in cells (Zanic et al., 2013). Surprisingly, although increasing growth rates by tubulin alone *in vitro* is accompanied by low catastrophe frequency, the significant increase in microtubule growth rate with XMAP215 was not accompanied by a suppression of catastrophe (Vasquez et al., 1994)(Zanic et al., 2013). Importantly, the effect of XMAP215 on the size of the GTP-cap is not known.

In this study, we investigate how XMAP215-promoted microtubule growth can simultaneously be fast and highly dynamic, displaying frequent microtubule catastrophes. First, we directly show that increasing tubulin concentration in the presence of EB1 increases microtubule growth rate and EB1 comet size, while simultaneously suppressing microtubule catastrophe frequency. Next, we add XMAP215 and demonstrate that XMAP215-driven increase in microtubule growth rate is accompanied by an increase in catastrophe frequency, as well as by an increase in EB1 comet size and brightness. Thus, the XMAP215-driven increase in catastrophe frequency is not a consequence of the GTP-cap size reduction. Rather, we demonstrate that XMAP215 increases microtubule growth fluctuations and induces tapered and curled microtubule ends. Our results suggest that XMAP215-induced destabilization of the growing microtubule end ultimately promotes microtubule catastrophe.

## RESULTS AND DISCUSSION

### Increasing the microtubule growth rate by increasing tubulin concentration correlates with an increase in GTP-cap size and suppression of microtubule catastrophe

To directly investigate the relationship between microtubule growth rate, catastrophe frequency, and GTP-cap size, we used an established assay for reconstitution of microtubule dynamics *in vitro* (Gell et al., 2010). Dynamic microtubule extensions were polymerized from GMPCPP-stabilized microtubule seeds using a range of tubulin concentrations (12 - 60 μM), and imaged with total internal reflection fluorescence (TIRF) microscopy (Figure 1A). The parameters of microtubule dynamics were determined using kymograph analysis (Zanic, 2016). To determine the size of the GTP-cap, we included 200 nM EB1-GFP in all conditions, and measured the intensity of EB1 comets at growing microtubule ends over a range of microtubule growth rates. The increase in microtubule growth rate achieved with tubulin titration was accompanied by a simultaneous suppression of microtubule catastrophe frequency (Figure 1B), consistent with studies using tubulin alone (Walker et al., 1988). In addition, increasing microtubule growth rates resulted in a linear increase in the average EB1-comet intensity (Figure 1C), consistent with previous reports (Bieling et al., 2007). Thus, our measurements directly establish an inverse correlation between the GTP-cap size and the microtubule catastrophe frequency, when increase in growth rate is achieved by increasing tubulin concentration in the presence of EB1 (Figure 1D). This finding is consistent with a model in which faster microtubule growth leads to a larger GTP-cap, which in turn provides enhanced protection against microtubule catastrophe.

**Figure 1.**
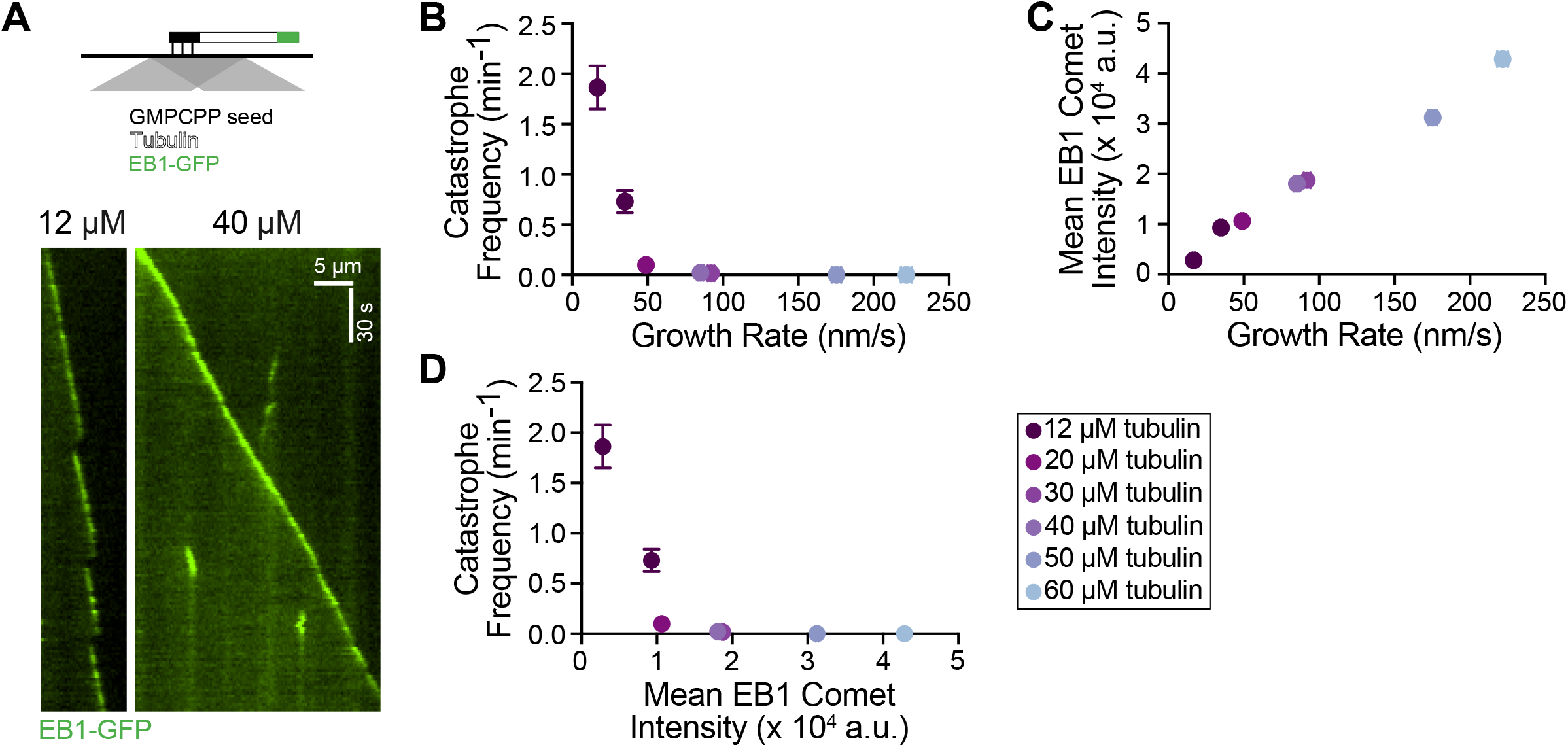
Increasing the microtubule growth rate by increasing tubulin concentration correlates with larger mean EB1 comets and suppression of microtubule catastrophe. (A) Top: schematic of TIRF assay. Dynamic microtubule extensions are polymerized from GMPCPP stabilized seeds using unlabeled tubulin in the presence of EB1-GFP. Bottom: Representative kymographs of microtubule plus ends grown with either 12 or 40 μM tubulin and 200 nM EB1-GFP. (B) Microtubule growth rate as a function of catastrophe frequency. (C) Microtubule growth rate replotted as a function of mean EB1 comet intensity. (D) Mean EB1 comet intensity replotted as a function of catastrophe frequency. 20 microtubules were analyzed per condition. Error bars are SEM.

### Increasing the microtubule growth rate using XMAP215 results in a simultaneous increase in microtubule catastrophe frequency

In cells, fast microtubule growth rates are achieved through the action of polymerases and other MAPs, including XMAP215 and EB1 (Akhmanova and Steinmetz, 2015). Interestingly, previous *in vitro* studies with XMAP215, either alone or in combination with EB1, reported that XMAP215-mediated increase in growth rate was not accompanied by a suppression of microtubule catastrophe frequency (Zanic et al., 2013; Vasquez et al., 1994). To investigate the relationship between microtubule catastrophe and microtubule growth rate in the presence of XMAP215, we quantified microtubule dynamics over a range of XMAP215 concentrations (3.13 - 200 nM) in the background of 20 μM tubulin and 200 nM EB1-GFP (Figure 2A). As expected, the microtubule growth rate increased as a function of XMAP215 concentration (Figure 2B). The increase in microtubule growth rate was accompanied by more frequent microtubule catastrophe events, even with the lowest XMAP215 concentration used (Figure 2C). This relationship between microtubule growth rate and catastrophe frequency in the presence of XMAP215 is in stark contrast to the one observed when growth rates were increased using tubulin titration (Figure 2D). Notably, XMAP215 led to a simultaneous increase in both growth rate and catastrophe frequency even in the absence of EB1 (Figure S1), demonstrating that the observed increase in catastrophe frequency can be directly attributed to XMAP215.

**Figure 2.**
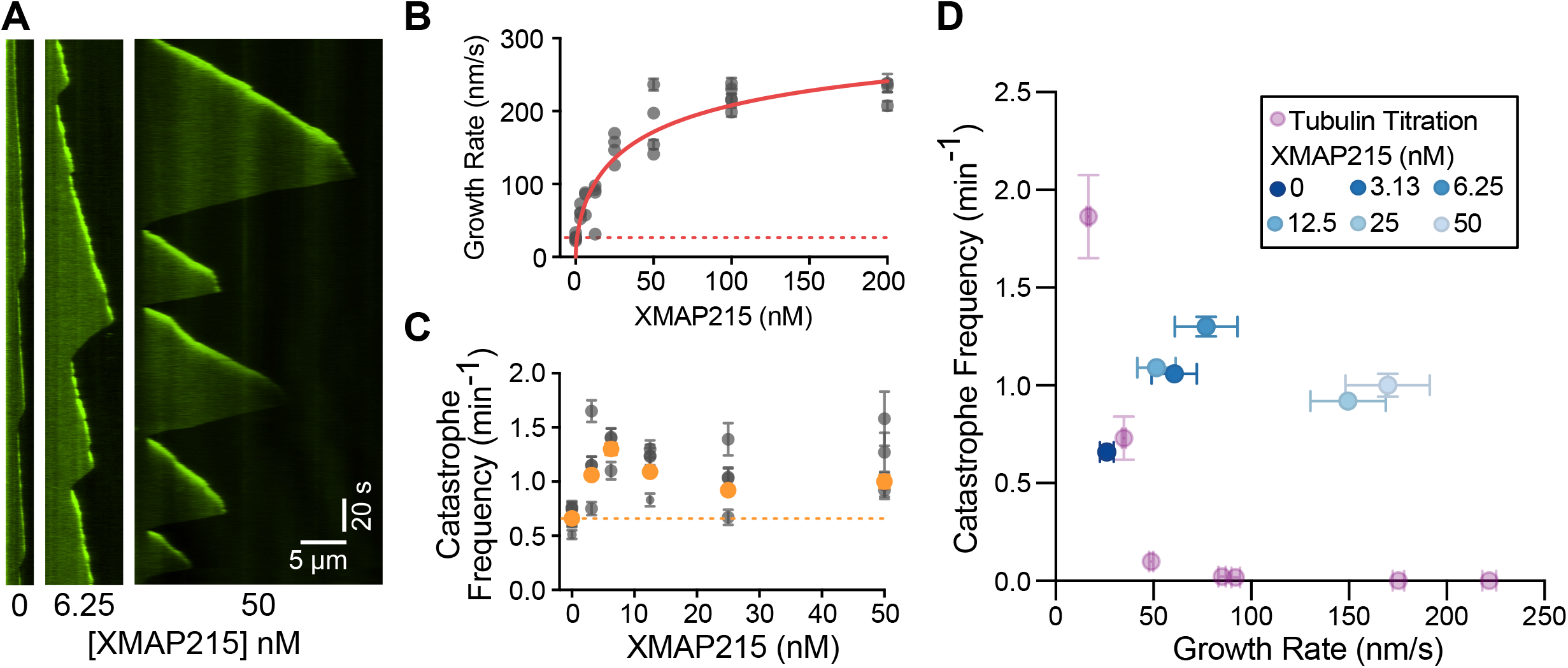
XMAP215 simultaneously increases microtubule growth rate and catastrophe frequency in the presence of EB1. (A) Representative kymographs of microtubule plus ends grown with 20 μM tubulin, 200 nM EB1-GFP, and corresponding amount of XMAP215 (nM). EB1-GFP is shown. Quantification of (B) microtubule growth rate and (C) catastrophe frequency as a function of XMAP215 concentration in the presence of 20 μM tubulin and 200 nM EB1-GFP. Error bars represent SEM. Each point represents 20 kymographs from one experimental repeat. The number of repeats per concentration were: 6, 4, 3, 4, 4, 4, 3. Dotted lines indicate the average values for the control (0 nM XMAP215). Solid red line in (B) is the data fit to the Hill equation. Orange points in (C) are the weighted averages for each condition. (D) Microtubule catastrophe frequency replotted as a function of microtubule growth rate for the XMAP215 titration along with the tubulin titration (Figure 1B).

### Promotion of catastrophe by XMAP215 is not achieved through a reduction in the GTP-cap size

One possible explanation for the observed increase in catastrophe frequency is that XMAP215 may be directly reducing the size of the protective GTP-cap at the growing microtubule end. While a linear increase in GTP-cap size with microtubule growth rate is well established for the tubulin titration (Figure 1C)(Bieling et al., 2007)(Strothman et al., 2019), whether the GTP-cap size increases when growth rate is increased by XMAP215 is not known. Our measurements of EB1 intensity with XMAP215 titration revealed a direct correlation between growth rate and EB1 intensity (Figure 3A). This finding suggests that increasing microtubule growth rate by XMAP215 also results in a larger GTP-cap size, similar to what was observed when the growth rate was increased using higher tubulin concentrations (Figure 1C).

**Figure 3.**
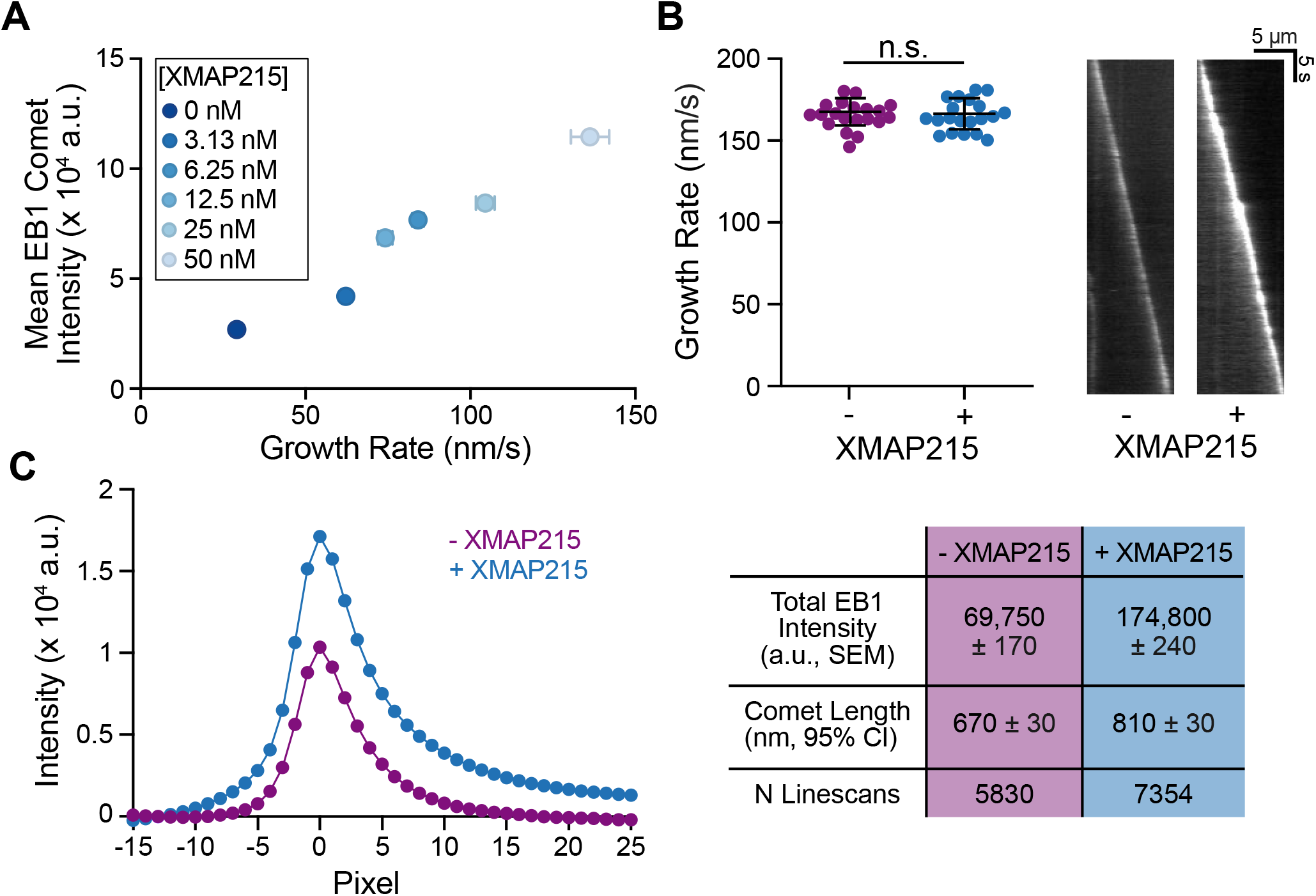
XMAP215 does not decrease the GTP-cap size. (A) Mean EB1 comet intensity is plotted against microtubule growth rate over a range of XMAP215 concentrations in the presence of 20 μM tubulin and 200 nM EB1-GFP. To generate each point, 20 microtubules were analyzed from one experiment, all experiments were performed on the same day. Error bars are SD. (B) Left: growth rates of microtubules polymerized with either 60 μM tubulin, 200 nM EB1-GFP (- XMAP215), or 20 μM tubulin, 200 nM EB1-GFP, and 25 nM XMAP215 (+ XMAP215). 20 growth events per condition with no significant difference in growth rate were analyzed, determined by unpaired t test. Means and SD are shown. Right: representative kymographs of EB1 localization in each condition. (C) The average EB1 comet profiles (left) and quantification (right) from 20 growth events per each condition.

To directly compare the mean GTP-cap size of microtubules grown in the presence or absence of XMAP215, we next performed growth-rate-matching experiments. We found that growth rates achieved with 60 μM tubulin and 200 nM EB1-GFP could be matched using 20 μM tubulin, 200 nM EB1-GFP and 12.5-25 nM XMAP215 (Figure 3B). To precisely compare the EB1 comet sizes, we generated averaged comet intensity profiles for each of the two conditions, following a previously established approach (see Methods)(Bieling et al., 2007). Surprisingly, we found that both the total intensity and the decay length of the EB1 comets were larger in the presence of XMAP215 (Figure 3C), in spite of the significantly higher catastrophe frequency in the XMAP215 condition when compared to the tubulin control (0.32 ± 0.04 min^−1^, SEM, N=57 and 0.005 ± 0.005 min^−1^, SEM, N=0 in the presence and absence of XMAP215, respectively). This finding directly demonstrates that promotion of microtubule catastrophe by XMAP215 is not a result of a decrease in the mean GTP-cap size.

### XMAP215 increases growth rate fluctuations and induces tapered microtubule ends

Our growth-rate-matching experiments provided an excellent dataset for a direct comparison of microtubule growth characteristics in the presence and absence of XMAP215. While the mean growth rates in two conditions were matched, we wondered whether the fluctuations in microtubule growth rate may differ between the two conditions. To investigate this possibility, we tracked microtubule growth and determined deviations from the mean velocity using linear regression (Figure 4A). We found that the sum of squared residuals (SSR) was significantly higher in the presence of XMAP215 (0.02 ± 0.01 μm^2^/s, mean ± SD, N=90) than in the tubulin control conditions (0.013 ± 0.007 μm^2^/s, mean ± SD, N=103) (Figure 4A), despite no difference in the mean growth rate (Figure S2A). This result was further corroborated by the mean squared displacement (MSD) analysis of the growing microtubule end positions in the presence and absence of XMAP215 (Figure S2B). Accompanying the increase in growth rate fluctuations, we additionally saw increased fluctuations in EB1 intensity in the presence of XMAP215 (Figure 4B), consistent with a previous study correlating the fluctuations in EB intensity with microtubule growth rate (Rickman et al., 2017). Thus, we conclude that microtubules polymerizing with XMAP215 display a higher degree of growth rate variability than those polymerizing at the same growth rates in the absence of XMAP215.

**Figure 4.**
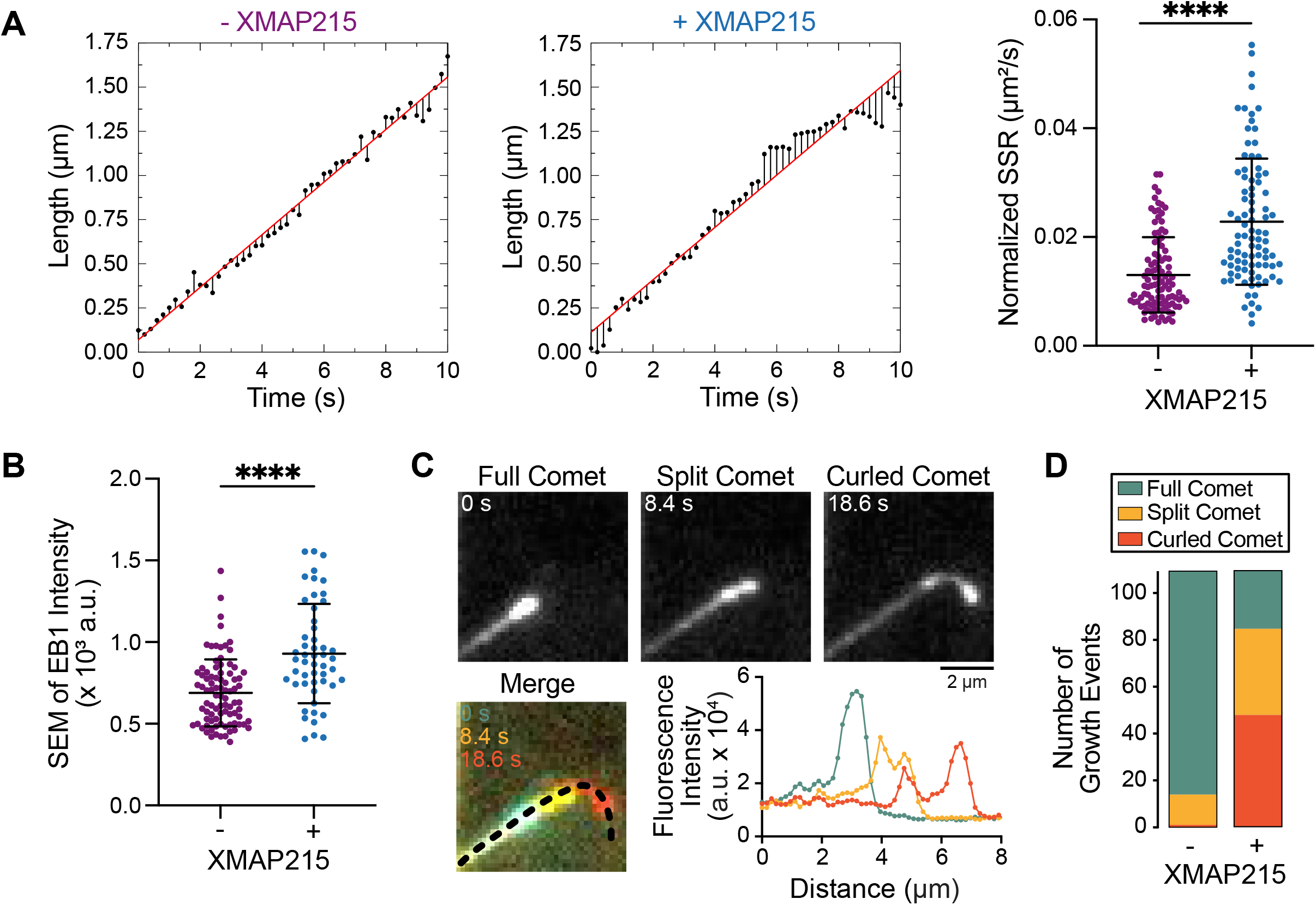
XMAP215 promotes fluctuations in microtubule growth and tapered microtubule ends. Growth-rate-matching conditions were achieved by either polymerizing microtubules with 60 μM tubulin, 200 nM EB1GFP (- XMAP215), or with 20 μM tubulin, 200 nM EB1GFP, and 12.5/25 nM XMAP215 (+ XMAP215). (A) Sum of squared residuals was determined from 10 s tracks that only displayed full comets. Left and center: representative example tracks from each condition are shown with microtubule tip position (black points) and residual for each time point (black lines), and linear regression to tip position (red line). Right: quantification of - XMAP215: 0.013 ± 0.007 μm^2^/s (mean ± SD, N = 103), and + XMAP215: 0.023 ± 0.011 μm^2^/s (mean ± SD, N = 90). ****, p < 0.0001 using unpaired Welch’s t test. (B) SEM of time-averaged EB1 intensities from 10-second segments tracks. - XMAP215: 0.7 ± 0.3 × 10^4^a.u. (mean ± SD, N = 87), and + XMAP215: 1.1 ± 0.04 x10^4^a.u. (mean ± SD, N = 51). ****, p < 0.0001 using unpaired Welch’s t test. A subset of tracks from (A, right) that displayed the same mean EB intensity was used. (C) EB1 comet morphology changes over time are classified into three different groups: full, split, and curled comets. Linescans of the growth trajectory over time reveal EB1 intensity changes that define the morphologies. (D) 110 microtubule growth events were tracked for no more than 2 min and classified to be the most disrupted end morphology observed. Number of growth events classified as either a full, split, or curled comet for each condition: - XMAP215: 96, 13, 1, and + XMAP215 25, 37, 48.

Interestingly, our high spatiotemporal-resolution tracking of EB1-GFP localization at microtubule ends polymerized with XMAP215 also revealed a range of comet morphologies over time (Movie and Figure 4C). Canonical EB localization has been previously described as a single peak of fluorescence that exponentially decays away from the direction of microtubule growth (Bieling et al., 2007), hereafter referred to as a ‘full comet’ (Figure 4C, left). However, in the presence of XMAP215, we observed frequent incidences of EB1 comets which appeared to split into two distinct intensity peaks, displaying a leading and a lagging comet, both of which were still growing in the original direction of growth (Figure 4C, center). Subsequent to comet splitting, we occasionally observed the lagging comet catching up to the leading comet, a phenomenon previously termed a ‘tip repair’ event (Aher et al., 2018)(Doodhi et al., 2016). Furthermore, we observed that a large number of split comet events led to a ‘curled comet’ morphology (Figure 4C, right). Curled comets were defined by the leading comet growing away from the original growth direction, resulting in the leading part of the polymer to bend. Quantification of the comet morphologies in the growth-rate-matching experiments revealed that microtubules polymerized with XMAP215 were six times more likely to display a tapered end (either split or curled comet) when compared to those grown at the same growth rate without XMAP215 (increase from 14 in the absence, to 85 in the presence of XMAP215 out of 110 comets quantified for each condition, Figure 4D). Given that the growth rates were the same between the control and XMAP215 conditions, these observations suggest that the increase in the frequency of tapered microtubule ends is a direct consequence of XMAP215.

### Microtubules grown with XMAP215 undergo catastrophe at faster growth rates and with more EB1 remaining

Our results thus far suggest that XMAP215 disrupts the structural integrity of the GTP-cap by inducing fluctuations in microtubule growth and promoting tapered microtubule ends. We hypothesized that these disruptions make microtubules more prone to catastrophe. To gain insight into the process of GTP-cap loss leading to catastrophe, we compared microtubule end position and EB1 intensity during catastrophe events using 0 nM and 25 nM XMAP215 conditions (in the background of 12 μM tubulin and 200 nM EB1-GFP), which both displayed robust catastrophe frequencies. We observed that microtubules polymerized in the absence of XMAP215 experienced a slowdown in growth rate prior to the onset of catastrophe, accompanied by a decrease in EB1 intensity (Figure 5A top). Previous work investigating tubulin-dilution-induced catastrophe found that a minimum of 29% occupancy in EB1 binding sites was necessary for protection against catastrophe (Duellberg et al., 2016). Similarly, we found that for spontaneously-occurring catastrophe in the absence of XMAP215, 25 ± 7% (mean ± SD, N=26) of EB1 binding sites remained occupied at the time of catastrophe (Figure 5B, Figure S3). In contrast, although microtubules polymerized in the presence of XMAP215 also experienced a slowdown prior to catastrophe (Figure 5A bottom), the switch to catastrophe occurred at comparatively faster growth rates (Figure 5C). Additionally, in the presence of XMAP215, microtubules underwent catastrophe with 38 ± 17% (mean ± SD, N=37) of EB1 binding sites occupied (Figure 5B), an amount significantly greater than in the control condition (p<0.001, Welch’s unpaired t-test). Our results demonstrate that microtubule ends grown with XMAP215 are inherently less stable, as they undergo catastrophe at faster growth rates and with more EB1, when compared to microtubules polymerized without XMAP215.

**Figure 5.**
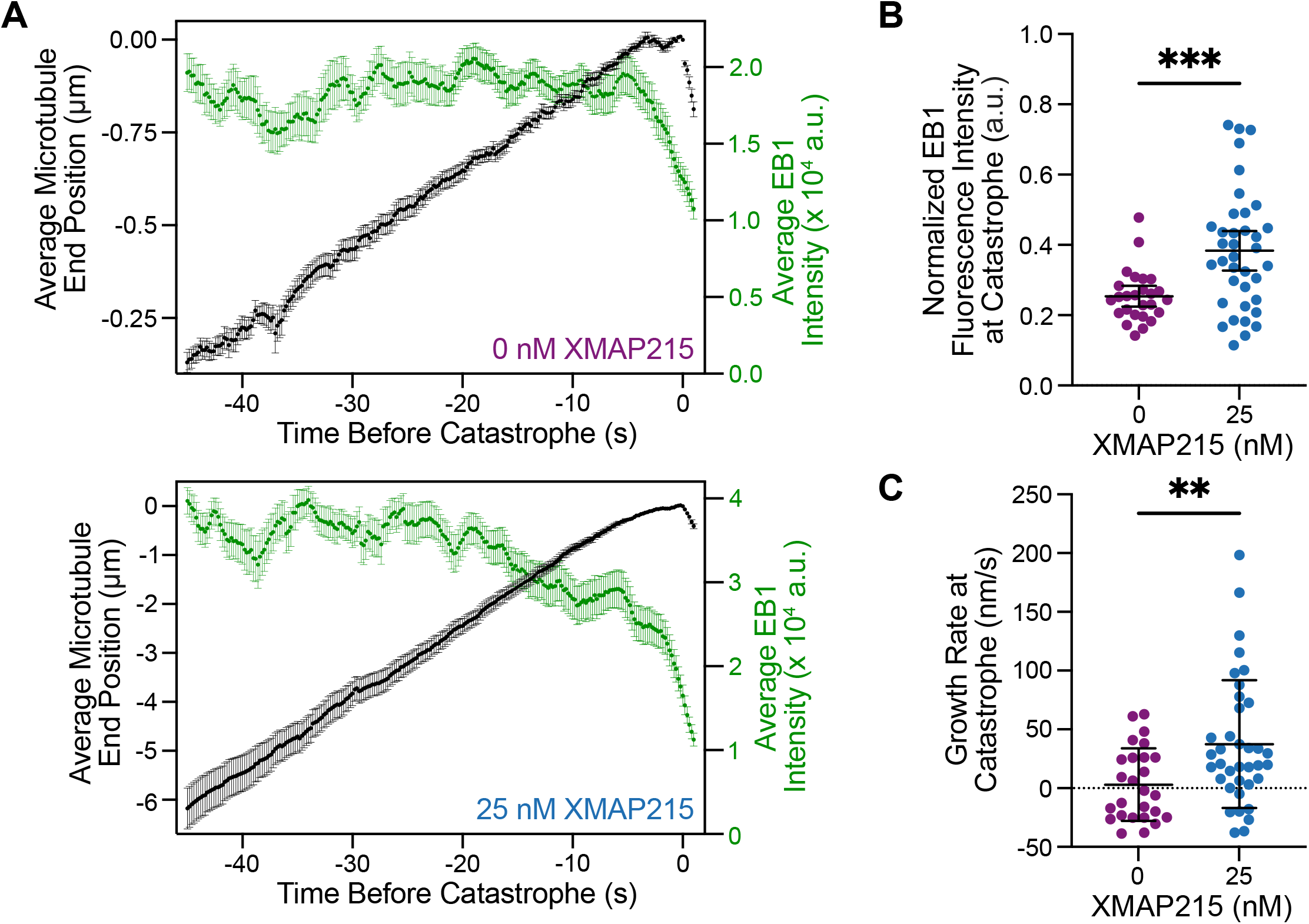
Microtubules grown in the presence of XMAP215 undergo catastrophe at faster growth rates and with more EB1. (A) Average microtubule end position and EB1 intensity over time. For 0 nM XMAP215, 30 tracks were averaged along their entire lifetime, with a minimum of 22 tracks for each time point. For 25 nM XMAP215, 38 tracks were averaged along their entire lifetime, with a minimum of 25 tracks for each time point. Average EB1 intensity was determined using a 5-frame (1 s) sliding window for each time point. Error bars represent SEM for both position and EB1 intensity. (B) EB1 intensity at the time of catastrophe was determined using 1 s prior to catastrophe (5 frames). EB1 fluorescence was normalized to the maximum intensity measured across all time points, independent of the condition (Fig. S3). 0 nM XMAP215 undergo catastrophe with 0.25 ± 0.07 a.u. (mean ± SD, N = 26), while 25 nM XMAP215 undergo catastrophe with 0.38 ± 0.17 a.u. (mean ± SD, N = 37). ***, p < 0.0001 using unpaired Welch’s t test. (C) Microtubule growth rate at the time of catastrophe was determined using 1 s prior to catastrophe (5 frames). 0 nM XMAP215: 3 ± 31 nm/s (mean ± SD, N = 26), and 25 nM XMAP215: 37 ± 54 nm/s (mean ± SD, N = 37). **, p = 0.0045, unpaired Welch’s t test.

## Conclusions

A cap of GTP-tubulin at the end of a growing microtubule is widely accepted as the determinant of microtubule stability (Mitchison and Kirschner, 1984)(Drechsel and Kirschner, 1994)(Desai and Mitchison, 1997)(Duellberg et al., 2016)(Roostalu et al., 2020). The size of the GTP-cap is defined by the balance between the addition of new GTP-tubulin dimers to the growing microtubule end, and the hydrolysis of GTP to GDP within the microtubule polymer. On its own, an increase in growth rate is expected to increase the size of the GTP-cap and thus confer enhanced stability to the growing microtubule. Indeed, our results using tubulin titration confirm that increase in growth rate is accompanied by an increase in EB1 comet size, as well as suppression of microtubule catastrophe. However, these findings raise the puzzling question of how simultaneously fast, yet highly dynamic microtubule growth, as observed in cells, can be achieved. One possibility to limit the size of the GTP-cap, and thus presumably facilitate catastrophe, is through the acceleration of the GTP-hydrolysis rate. This mechanism has been proposed for EB proteins, which promote microtubule catastrophe even while inducing a slight increase in growth rate (Bieling et al., 2007)(Zhang et al., 2015)(Vitre et al., 2008). In contrast, our results demonstrate that XMAP215 simultaneously promotes microtubule growth and catastrophe frequency without accelerating the GTP-hydrolysis rate, or otherwise decreasing the mean GTP-cap size. In fact, we find that the mean size of the EB comets is even larger when microtubule growth acceleration is achieved through the action of XMAP215.

Aside from the nucleotide composition, both kinetics of microtubule assembly and the structure of the growing microtubule end remain areas of intense interest (Kerssemakers et al., 2006) (Schek et al., 2007)(Gardner et al., 2011a)(Mickolajczyk et al., 2019)(Guesdon et al., 2016)(Estévez-Gallego et al., 2020)(Mcintosh et al., 2018)(Atherton et al., 2018)(Gudimchuk et al., 2020). The addition of inherently curved GTP-tubulin dimers to the ends of microtubule protofilaments and their subsequent straightening into closed polymer lattice have been reported to result in a variety of intermediate structures, including disconnected and splayed individual protofilaments (Mcintosh et al., 2018)(Gudimchuk et al., 2020), multi-protofilament sheets (Chrétien et al., 1995)(Guesdon et al., 2016)(Atherton et al., 2018) and overall tapered microtubule ends (Mandelkow et al., 1991)(Reid et al., 2019). Microtubule end structures can vary with tubulin from different species (Orbach and Howard, 2019), and can be further modulated by microtubule-associated-proteins and drugs (Chen and Hancock, 2015)(Chen et al., 2019)(Aher et al., 2018)(Doodhi et al., 2016)(Arnal et al., 2000). In the case of XMAP215, the acceleration of microtubule growth has been linked to its ability to bind curved tubulin dimers both in solution and at the microtubule end (Ayaz et al., 2012)(Brouhard and Rice, 2014). XMAP215 is thought to stabilize an intermediate state in microtubule assembly, effectively acting as an enzyme activator for the polymerization reaction. Such a catalytic mechanism can explain how XMAP215 greatly accelerates microtubule polymerization in the presence of soluble tubulin, but also promotes depolymerization of stabilized microtubules in the absence of soluble tubulin (Brouhard et al., 2008). Given that XMAP215 was reported to act as a processive polymerase, with each XMAP215 molecule promoting addition of ~25 tubulin dimers (Brouhard et al., 2008), we speculate that XMAP215 molecules primarily drive elongation of individual protofilaments, resulting in less coordinated protofilament growth. Indeed, our observations of EB1 comet splitting and curling suggest that polymerization is not synchronized among all protofilaments. Instead, some protofilaments grow faster while others grow slower, to produce an overall tapered microtubule ends. Given that EBs localize to the interface of four tubulin dimers (Maurer et al., 2012), our observation of leading comets suggests the presence of multiple laterally-connected protofilaments within these protrusions. The existence of tapered and open microtubule ends can further facilitate EB1 targeting (Reid et al., 2019), consistent with our observations of brighter EB1 comet intensities on XMAP215-grown microtubule ends. Importantly, although we used EB1 to visualize the nucleotide composition and structure of growing microtubule ends, our observation of XMAP215-dependent promotion of catastrophe in the absence of EB1 demonstrates that XMAP215 on its own, rather than through enhanced targeting of EB1, promotes microtubule catastrophe.

Microtubule catastrophe is a complex phenomenon known not to follow first-order kinetics; rather, a microtubule aging process ultimately leads to rapid microtubule disassembly (Odde et al., 1995)(Gardner et al., 2013)(Gardner et al., 2011b). Previous studies implicated evolution of the microtubule-end structure and accumulation of structural defects as steps preceding microtubule catastrophe (Gardner et al., 2011b)(Coombes et al., 2013)(Bowne-Anderson et al., 2013). Our results demonstrate that XMAP215, in addition to inducing dramatic microtubule end morphologies, promotes large fluctuations in growth and comet intensities even when microtubule ends display ‘full’ comets. We speculate that XMAP215 induces ‘sloppy’ microtubule growth, as both the structure and the nucleotide composition of the growing microtubule end are highly dynamic in the presence of XMAP215. Notably, we find that even at the moment of catastrophe, microtubule ends polymerized with XMAP215 display faster growth rates and more EB1 localization, compared to microtubules grown without XMAP215. Our results reveal that the switch to catastrophe can occur with GTP-cap densities significantly higher than the threshold previously established by tubulin-dilution studies (Duellberg et al., 2016), suggesting that structural changes induced by XMAP215 can override the protective effects of the nucleotide cap.

Finally, while the polymerase effects of XMAP215 are dose-dependent, such that the maximum microtubule growth promotion is reached in the ~100 nM range, we find that XMAP215’s promotion of microtubule catastrophe reaches its full effect even at the lowest concentration of XMAP215 tested (3.13 nM). This observation provides additional evidence that the processes of microtubule growth and catastrophe are inherently decoupled. Future structural studies and the refinement of existing computational models (Bowne-Anderson et al., 2013)(VanBuren et al., 2002)(VanBuren et al., 2005)(Margolin et al., 2012)(Zakharov et al., 2015)(Kim and Rice, 2019)(Michaels et al., 2020)(Gudimchuk et al., 2020) will be necessary to unravel the full complexity of microtubule dynamics. Notwithstanding, the ability to independently control microtubule growth and catastrophe is at the very core of microtubule regulation in cells, enabling the complex, dynamic remodeling of the microtubule cytoskeleton.

## ACKNOWLEDGEMENTS

We thank W. Hancock, R. Ohi, the members of the Zanic laboratory and Vanderbilt Microtubules and Motors Club for discussions and feedback. XMAP215 construct was a kind gift from G. Brouhard (McGill University, Montreal, CA).

This work was supported by National Institutes of Health grant R35GM119552 to M. Zanic. V. Farmer acknowledges support from National Institutes of Health grant T32GM008320 and American Heart Association Predoctoral Fellowship 19PRE34380083. M. Zanic acknowledges support from the Human Frontier Science Program and the Searle Scholars Program.

The authors declare no competing financial interests.

## AUTHOR CONTRIBUTIONS

MZ, VF, and GA conceptualized the project, designed the research, and wrote the manuscript. SH and VF contributed reagents. VF performed experiments. GA developed image analysis scripts. VF and GA performed image and data analysis.

## MATERIALS AND METHODS

### Protein preparation

Bovine brain tubulin was purified as previously described through cycles of polymerization and depolymerization in a high molarity PIPES buffer (Castoldi and Popov, 2003). Tubulin was labeled with either TAMRA or Alexa 647 (Invitrogen) as previously described (Hyman et al., 1991). For imaging purposes, labeled tubulin was used at a ratio of 10% of the final tubulin concentration. XMAP215-7his expression construct was a kind gift from G. Brouhard, McGill University, Montreal, QC, Canada. XMAP215 was expressed in Sf9 cells using the Bac-to-Bac system (Invitrogen) and purified using a HisTrap followed by gel filtration (adapted from Brouhard et al., 2008), and stored in 10 mM Bis-Tris, 10 mM TrisHCl, 100 mM KCl, 1 mM DTT, 10% glycerol, pH 6.6. EB1-GFP was expressed in *Escherichia coli* and purified as previously described (Zanic et al., 2009), and stored in 10 mM Bis-Tris, 10 mM TrisHCl, 100 mM KCl, 1 mM DTT, 5% glycerol, pH 6.6. Protein concentration was determined using absorbance at λ = 280 nm.

### TIRF microscopy

Imaging was performed using a Nikon Eclipse Ti microscope with a 100X/1.49 n.a. TIRF objective; Andor iXon Ultra EM-CCD camera; 488-, 561-, and 640-nm solid-state lasers (Nikon Lu-NA); Finger Lakes Instruments HS-625 high speed emission filter wheel; and standard filter sets. A Tokai Hit objective heater was used to maintain the sample at 35°C. Samples were imaged in chambers constructed as previously described (Gell et al., 2010). In brief, three strips of Parafilm were sandwiched between a 22 × 22 mm and 18 × 18 mm silanized coverslips to create two narrow channels for the exchange of reaction solution. The channel surface was treated with 0.02 μg/μl anti-TAMRA antibody (Invitrogen) followed by 1% Pluronic F127 (Sigma) prior to use. Images were acquired using NIS-Elements (Nikon).

### Assay conditions

GMPCPP-stabilized, 25% TAMRA-labeled microtubules were polymerized as previously described (Hunter et al., 2003) and immobilized to coverslips using anti-TAMRA antibody (Gell et al., 2010). The imaging buffer consisted of BRB80 supplemented with 40 mM glucose, 40 μg/ml glucose oxidase, 25 μg/ml catalase, 0.08 mg/ml casein, 10 mM DTT, and 0.1% methylcellulose. Reactions containing imaging buffer, concentrations of tubulin ranging from 12 to 60 μM tubulin, 1 mM GTP, and proteins at the concentrations indicated in the text were introduced into the imaging chamber. XMAP215 storage buffer was consistently kept at a final concentration of 4X-dilution (2.5 mM Bis-Tris, 2.5 mM TrisHCl, 25 mM KCl, 250 nM DTT, 2.5% glycerol) for the entire XMAP215 titration.

### Microtubule dynamics analysis

Quantification of microtubule dynamics parameters was performed using 5-pixel width kymographs of the tubulin channel generated in Fiji (Schindelin et al., 2012) as described previously (Zanic, 2016). For each experiment, 20 kymographs were analyzed for dynamic parameter quantification. Microtubule polarity was determined by measuring growth rates of the two ends of a microtubule, with the faster-growing end being designated as the plus end and the slower-growing end the minus end. Catastrophe events were designated as a switch from growth to shrinkage that decreased microtubule length by more than 2 pixels (320 nm). Catastrophe frequency was calculated as the number of events divided by the total time spent in growth for an individual experiment, with an error of the square root of the number of events divided by the total time spent in growth (counting error).

### EB1 comet analysis (Figures 1–3)

EB1 comets were quantified using a series of custom MATLAB (version R2020a, MathWorks) scripts as described before (Strothman et al., 2019). Briefly, the beginning and end of individual growth events were manually determined from kymographs, and the initial estimate of microtubule tip position over time was obtained assuming a constant growth rate. For each time frame, the pixel with the brightest EB1 intensity within a window (± 5 or ± 10 pixels for – XMAP215 and + XMAP215 conditions, respectively) around the estimated tip position was assigned as the microtubule tip position. For an entire microtubule growth event, the average tip intensity and its standard deviation were calculated. The temporal frames with tip intensities lower than one standard deviation away from the mean were eliminated. The remaining tip positions were subsequently fit by linear regression, and mean and standard deviation of the fit residuals were determined. The temporal frames with ± 1 standard deviation away from the mean of the residuals were eliminated, and the remaining positions were fit by linear regression to assign a growth rate to each event. The tip positions from the remaining temporal frames were aligned using the maximum EB intensity. The microtubule lattice intensity was determined by averaging the intensity 15 pixels away from the tip, which was subsequently subtracted from the intensity of all pixels along the intensity profile. For a given experimental condition, individual growth events were further excluded if their growth rate was not within ± 1 standard deviation from the mean growth rate. Remaining events were averaged to determine a super-averaged intensity profile, where error is the standard error of the mean.

To determine EB1 comet length, the super-averaged intensity profile was fit to an exponential decay convolved with a Gaussian function:

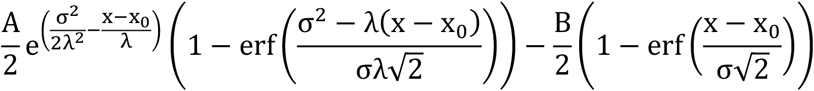

where *A* is the intensity value at the tip, *B* is the difference between average lattice intensity and solution background, *σ* is the experimentally-determined full width at half maximum of the point spread function, *x*_0_ is the offset in the tip position due to convolution, and *λ* is the comet decay length. To determine the total EB1 intensity along the super-averaged intensity profile, the pixels with positive intensity (i.e. greater than lattice background) were summed. The error for total EB1 intensity was determined using error-propagation.

### Determination of variability in microtubule growth (Figure 4)

Individual microtubule growth events from the growth-rate-matching conditions which displayed a full comet during their lifetime were subjected to automated tracking. Images were background-subtracted using an average 5-pixel rolling ball subtraction. The EB1 channel was tracked with FIESTA’s single particle tracker (Ruhnow et al., 2011) using MATLAB. Then, a custom MATLAB code was used to divide the output trajectories into continuous 10-second segments, allowing for gaps of no more than a total of 1-second within a given segment. The variations from the mean growth rate within the 10-s segments were quantified by performing residuals analysis as previously described (Lawrence et al., 2018). Briefly, using a custom MATLAB code, a linear function was fit to the length-versus-time data points to determine the mean growth rate. The sum of squared residuals (SSR) was calculated and normalized by the segment duration. For growth-rate matching experiments, only trajectories with mean growth speeds between 110 and 180 nm/s were considered. Outliers based on normalized SSR were identified using MATLAB function “isoutlier” and subsequently discarded. Unpaired t-test with Welch’s correction was used to determine p-values for mean velocity and normalized SSR between ±XMAP215 conditions. The same selected segments were subjected to Mean Squared Displacement analysis using MATLAB-based “msdanalyzer” (Tarantino et al., 2014). A quadratic function (Gardner et al., 2011a) was fit to the first 5 seconds of the MSD curve:

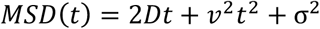

where *D* is diffusion coefficient, *v* is mean growth rate, σ is the positional error. The fit was weighted by the inverse of the standard deviation of the MSD curve determined by msdanalyzer.

### Determination of EB1 intensity (Figure 4)

To determine EB1 intensity at microtubule ends a custom MATLAB function was used. Microtubule end position was determined either by tubulin or EB1 signal for XMAP215 titration or growth-rate-matching experiments, respectively. For each temporal frame, the image was rotated centering around the end position, so that the microtubule lies horizontally. The brightest intensity value within 5-lattice-pixels and 1-solution-pixel was assigned as maximum EB1 intensity (5-pixel thickness, i.e. 5×6 pixel^2^ area). Solution background intensity was determined by shifting the 5×6 pixel^2^ area up and down 5 pixels, and the mean intensity was calculated. Temporal frames with <25 pixels available for background determination were discarded. For each temporal frame, the mean background intensity was then subtracted from the EB1 intensities. For the comparison of EB1 intensities during growth in growth-rate-matching conditions, time-averaged EB1 intensities were calculated using 10-second window size.

### Determination of microtubule end morphology in growth-rate matching experiments (Figure 4)

Microtubule dynamic assays were carried out in the presence of 200 nM EB1-GFP with either high tubulin concentration (60 μM) or low tubulin concentration (12 μM) and XMAP215 (12.5 or 25 nM). Imaging in the EB1 channel was acquired at 5 frames-per-second to allow for high spatiotemporal tracking of the microtubule ends. Individual microtubule growth events were tracked for up to 2 min and the average microtubule growth rate was determined for each growth event. Two sets of 110 growth events, one for each condition (-/+ XMAP215), with no significant difference in their growth rates were scored for catastrophe (as defined above) and end morphology. End morphology was classified into three categories based on EB1 signal at the growing microtubule end: full, split, or curled comet. Kymographs were produced from 7-pixel wide linescans (1120 nm) and subsequently used to determine if EB1 localized in one single peak at the end of a growing microtubule, classified as a ‘full’ comet. If two peaks of intensity could be resolved (>2 pixels) for more than 1 seconds (5 frames) the comet was considered to be ‘split’. A ‘curled’ comet was preceded by a splitting event with the leading comet having grown outside the 7-pixel-wide linescan.

### Determination of the velocity and EB1 intensity at catastrophe (Figure 5)

Individual microtubule growth events from the 0 or 25 nM XMAP215 conditions, which displayed only a full comet morphology during their lifetime, were subjected to automatic tracking. For each individual growth event, microtubule position was determined from the tubulin signal using TipTracker v3 (Prahl et al., 2014). First, both x- and y-coordinates of the microtubule end from each temporal frame, except the initial and final frame, were preprocessed to eliminate tracking noise. If the difference between coordinates of the current frame and the previous frame was larger than 1000 nm, the current coordinate value was eliminated and a new coordinate value was interpolated using the previous and subsequent frame, assuming a linear growth rate. To further minimize tracking noise, the “smoothdata” function in MATLAB was used with the “movmedian” method and a 5-frame (1-second) window size. The end position was determined using smoothened coordinates. Finally, the time of catastrophe was approximated manually and subsequently fine-tuned using the following automated analysis: each time point in the time interval of 10 frames before and after the manually-approximated time of catastrophe was assigned an instantaneous rate using a linear fit over a 3-frame sliding window. Then, starting from 8 frames after catastrophe and moving backwards in time, if three consecutive frames had velocity value greater than −50 nm/s, the latest of the three temporal frames was assigned as the time of catastrophe. After determining the time of catastrophe, the end-position of growth events over time were aligned to generate an averaged microtubule tip position using a custom MATLAB code. For each microtubule, time and position values were offset to assign catastrophe event to (0,0). Then, the mean and standard error of the mean for the positions at each time point over different growth events were calculated.

EB1 intensity was determined as described above, using a 1-second window size. The intensity error was determined by propagating the standard error of the mean of the solution background. After determining EB intensity as a function of time for each growth event, the intensities were averaged over all growth events at every time point, with error being the standard error of the mean, both weighted with the inverse of squared error of intensities. ROUT method with Q=1% identified 2 outlier points which were subsequently removed.

Microtubule growth events that underwent catastrophe were further evaluated to determine the growth rate prior to catastrophe. Briefly, a custom MATLAB function was used to perform a linear fit to the length-vs-time segments. To determine instantaneous growth rate at the time of catastrophe (T=0 seconds), a 1-second (5-frame) window size (i.e. from T=-1 seconds to T=0 seconds) was used. Growth rates preceding a catastrophe event were determined using 5-second intervals up to 45 seconds prior to catastrophe. Errors in growth rate were calculated as (CI_high_ – CI_low_)/2, where CI_high_ and CI_low_ are upper and lower 95% confidence intervals from the linear fits. ROUT method with Q=1% identified 2 outlier points which were subsequently removed.

## SUPPLEMENTAL MATERIAL

**Movie. EB1 comet morphologies at the end of a growing microtubules.** Time-lapse of a microtubule grown in the presence of 20 μM tubulin, 200 nM EB1-GFP, and 25 nM XMAP215. Scale bar is 2 μm.

**Figure S1.**
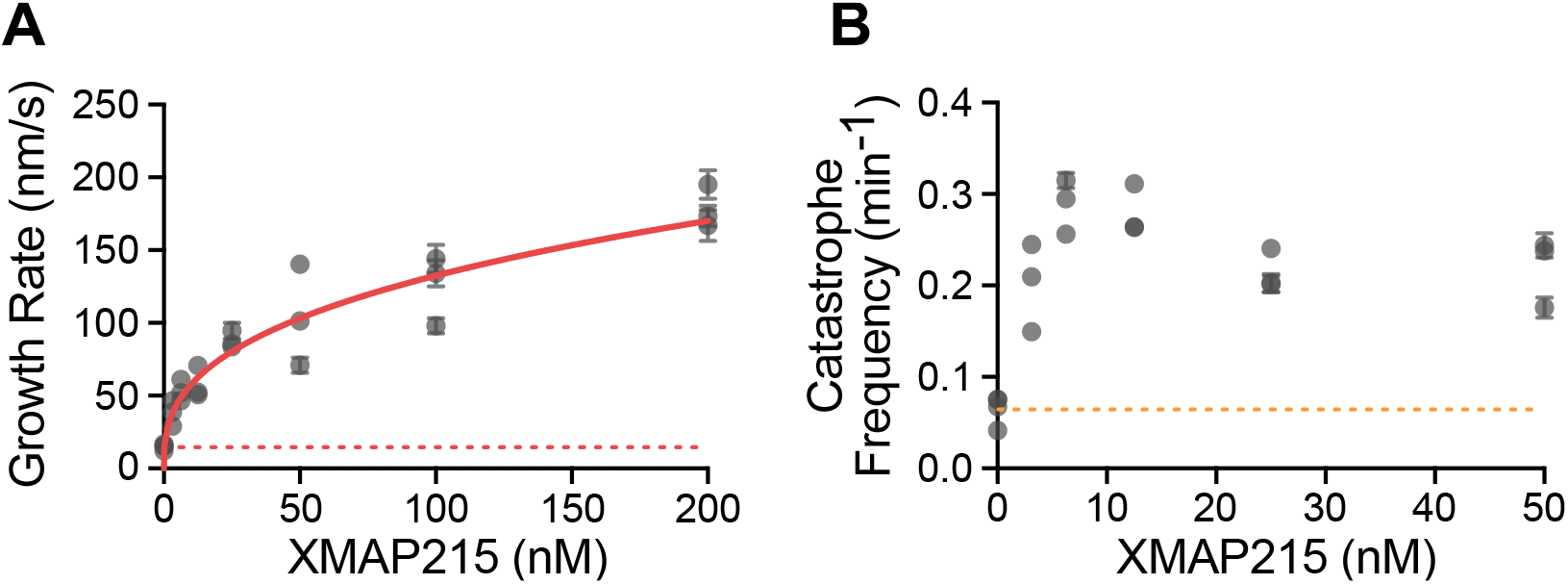
Simultaneous increase of microtubule growth rate and catastrophe frequency by XMAP215 occurs independently of EB1. Microtubule growth rate and catastrophe frequency were quantified over a titration of XMAP215 in the background of fixed tubulin concentration (20 μM). Each point represents 20 kymographs from one experimental repeat. The number of repeats per concentration were: 4, 3, 3, 3, 3, 3, 3. (A) Microtubule growth rate as a function of XMAP215 concentration. Dotted red line represents the growth rate in control conditions and solid red line is data fit to the Hill equation. Error bars are SEM. (B) Microtubule catastrophe frequency as a function of XMAP215 concentration. Dotted orange line represents the catastrophe frequency in control conditions. Error bars represent counting error.

**Figure S2.**
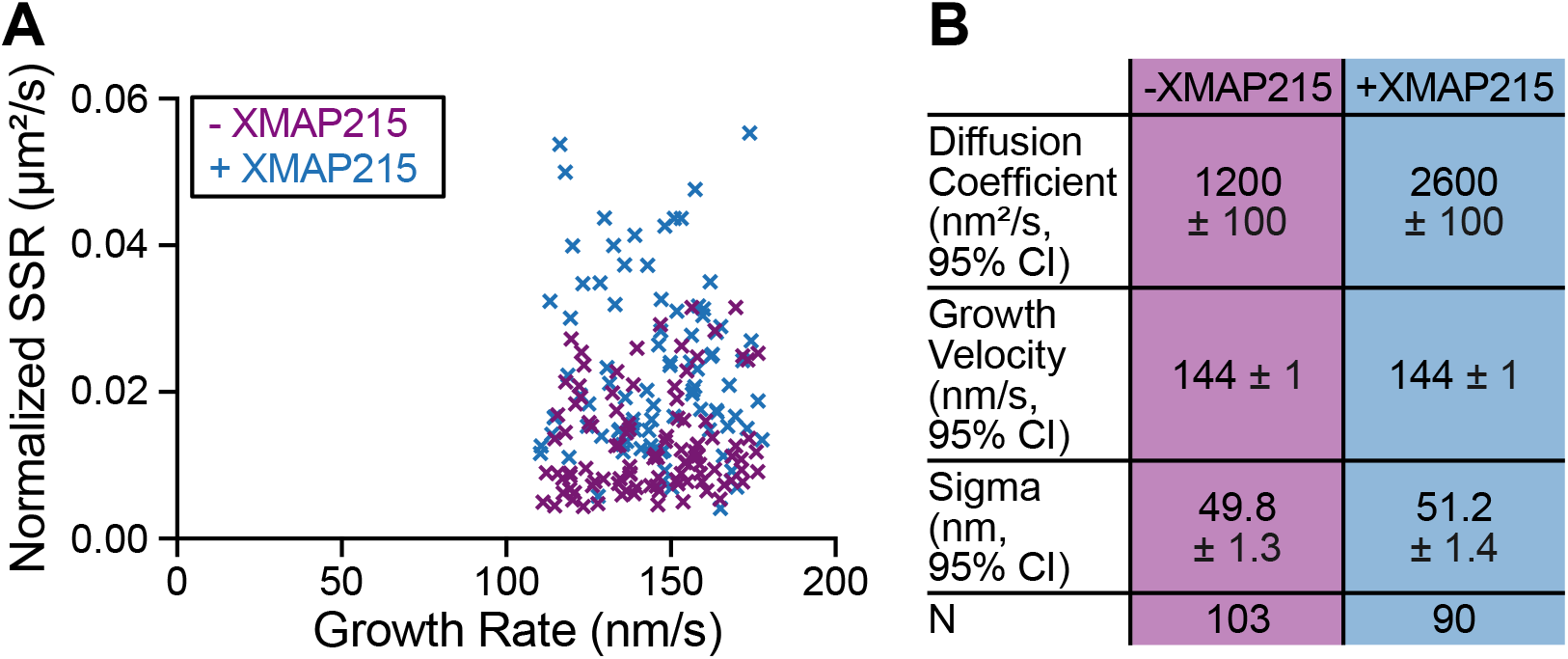
Growth rate matching conditions. (A) Growth events used for SSR analysis in Figure 4A were selected to have no significant difference in growth rate: - XMAP215; 143 ± 18 nm/s (mean ± SD, N = 103), and + XMAP215; 145 ± 18 nm/s (mean ± SD, N = 90). (B) Diffusion coefficients were determined by Mean Squared Displacement analysis by fitting a quadratic function (Gardner et al. 2011).

**Figure S3.**
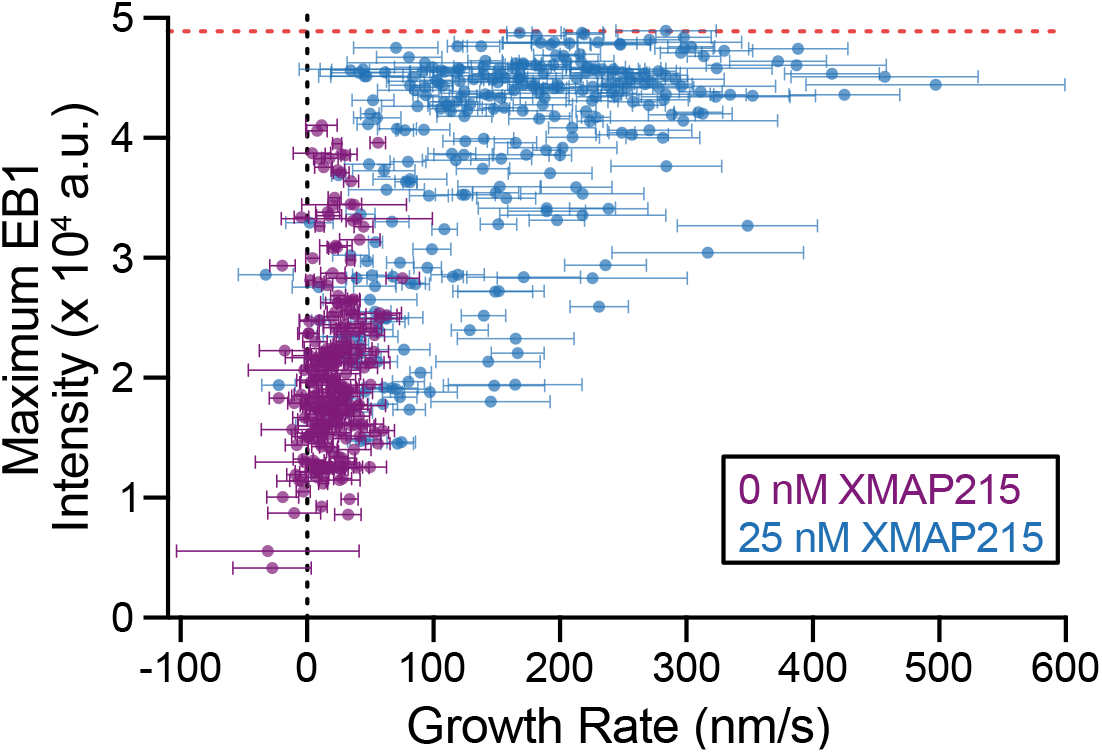
Maximum EB1 intensity positively scales with microtubule growth rate and plateaus, indicating saturation of binding sites. (A) Microtubule growth rate and EB1 intensity for every track were determined at 5 s intervals using a sliding window of either 5 s or 1 s, respectively. Error is 95% CI of the fit for growth rate and propagated error using SEM of background intensity. Dotted red line represents the saturating EB1 intensity, used for normalization in Figure 5B.

## REFERENCES

Aher, A., M. Kok, A. Sharma, A. Rai, N. Olieric, R. Rodriguez-Garcia, E.A. Katrukha, T. Weinert, V. Olieric, L.C. Kapitein, M.O. Steinmetz, M. Dogterom, and A. Akhmanova. 2018. CLASP Suppresses Microtubule Catastrophes through a Single TOG Domain. Dev. Cell. 46:40–58. doi:10.1016/j.devcel.2018.05.032.

Akhmanova, A., and M.O. Steinmetz. 2008. Tracking the ends: a dynamic protein network controls the fate of microtubule tips. Nat. Rev. Mol. Cell Biol. 9:309–322. doi:10.1038/nrm2369.

Akhmanova, A., and M.O. Steinmetz. 2015. Control of microtubule organization and dynamics: two ends in the limelight. Nat. Rev. Mol. Cell Biol. 16:711–26. doi:10.1038/nrm4084.

Al-Bassam, J., and F. Chang. 2011. Regulation of microtubule dynamics by TOG-domain proteins XMAP215/Dis1 and CLASP. Trends Cell Biol. 21:604–614. doi:10.1016/j.tcb.2011.06.007.

Arnal, I., E. Karsenti, and A.A. Hyman. 2000. Structural transitions at microtubule ends correlate with their dynamic properties in Xenopus egg extracts. J. Cell Biol. 149:767–774. doi:10.1083/jcb.149.4.767.

Atherton, J., M. Stouffer, F. Francis, and C.A. Moores. 2018. Microtubule architecture in vitro and in cells revealed by cryo-electron tomography. Acta Crystallogr. Sect. D Struct. Biol. 74:1–13. doi:10.1107/S2059798318001948.

Ayaz, P., X. Ye, P. Huddleston, C.A. Brautigam, and L.M. Rice. 2012. A TOG : αβ-tubulin Complex Structure Reveals Conformation-Based Mechanisms for a Microtubule Polymerase. Science (80-.). 337:857–60. doi:10.1126/science.1221698.

Bieling, P., L. Laan, H. Schek, E.L. Munteanu, L. Sandblad, M. Dogterom, D. Brunner, and T. Surrey. 2007. Reconstitution of a microtubule plus-end tracking system in vitro. Nature. 450:1100–1105. doi:10.1038/nature06386.

Bowne-Anderson, H., M. Zanic, M. Kauer, and J. Howard. 2013. Microtubule dynamic instability: A new model with coupled GTP hydrolysis and multistep catastrophe. BioEssays. 35:452–461. doi:10.1002/bies.201200131.

Brouhard, G.J., and L.M. Rice. 2014. The contribution of αβ-tubulin curvature to microtubule dynamics. J. Cell Biol. 207:323–334. doi:10.1083/jcb.201407095.

Brouhard, G.J., J.H. Stear, T.L. Noetzel, J. Al-Bassam, K. Kinoshita, S.C. Harrison, J. Howard, and A.A. Hyman. 2008. XMAP215 Is a Processive Microtubule Polymerase. Cell. 132:79–88. doi:10.1016/j.cell.2007.11.043.

Castoldi, M., and A. V. Popov. 2003. Purification of brain tubulin through two cycles of polymerization-depolymerization in a high-molarity buffer. Protein Expr. Purif. 32:83–88. doi:10.1016/S1046-5928(03)00218-3.

Chen, G.-Y., J.M. Cleary, A.B. Asenjo, Y. Chen, J.A. Mascaro, D.F.J. Arginteanu, H. Sosa, and W.O. Hancock. 2019. Kinesin-5 Promotes Microtubule Nucleation and Assembly by Stabilizing a Lattice-Competent Conformation of Tubulin. Curr. Biol. 29:2259–2269.e4. doi:10.1016/j.cub.2019.05.075.

Chen, Y., and W.O. Hancock. 2015. Kinesin-5 is a microtubule polymerase. Nat. Commun. 6:8160. doi:10.1038/ncomms9160.

Chrétien, D., S.D. Fuller, and E. Karsenti. 1995. Structure of growing microtubule ends: Two-dimensional sheets close into tubes at variable rates. J. Cell Biol. 129:1311–1328. doi:10.1083/jcb.129.5.1311.

Coombes, C.E., A. Yamamoto, M.R. Kenzie, D.J. Odde, and M.K. Gardner. 2013. Evolving tip structures can explain age-dependent microtubule catastrophe. Curr. Biol. 23:1342–1348. doi:10.1016/j.cub.2013.05.059.

Desai, A., and T.J. Mitchison. 1997. Microtubule Polymerization Dynamics. Annu. Rev. Cell Dev. Biol. 13:83–117. doi:10.1146/annurev.cellbio.13.1.83.

Doodhi, H., A.E. Prota, L.C. Kapitein, A. Akhmanova, and M.O. Steinmetz. 2016. Termination of Protofilament Elongation by Eribulin Induces Lattice Defects that Promote Microtubule Catastrophes. Curr. Biol. 26:1713–1721. doi:10.1016/j.cub.2016.04.053.

Drechsel, D.N., and M.W. Kirschner. 1994. The minimum GTP cap required to stabilize microtubules. Curr. Biol. 4:1053–1061. doi:10.1016/S0960-9822(00)00243-8.

Duellberg, C., N.I. Cade, D. Holmes, and T. Surrey. 2016. The size of the EB cap determines instantaneous microtubule stability. Elife. 5:1–23. doi:10.7554/eLife.13470.

Estévez-Gallego, J., F. Josa-Prado, S. Ku, R.M. Buey, F.A. Balaguer, A.E. Prota, D. Lucena-Agell, C. Kamma-Lorger, T. Yagi, H. Iwamoto, L. Duchesne, I. Barasoain, M.O. Steinmetz, D. Chrétien, S. Kamimura, J.F. Díaz, and M.A. Oliva. 2020. Structural model for differential cap maturation at growing microtubule ends. Elife. 9:1–26. doi:10.7554/eLife.50155.

Gard, D.L., B.E. Becker, and S. Josh Romney. 2004. MAPping the eukaryotic tree of life: Structure, function, and evolution of the MAP215/Dis1 family of microtubule-associated proteins. Int. Rev. Cytol. doi:10.1016/S0074-7696(04)39004-2.

Gard, D.L., and M.W. Kirschner. 1987. A microtubule-associated protein from Xenopus eggs that specifically promotes assembly at the plus-end. J. Cell Biol. 105:2203–2215. doi:10.1083/jcb.105.5.2203.

Gardner, M.K., B.D. Charlebois, I.M. Janosi, J. Howard, A.J. Hunt, and D.J. Odde. 2011a. Rapid microtubule self-assembly kinetics. Cell. 146:582–592. doi:10.1016/j.cell.2011.06.053.

Gardner, M.K., M. Zanic, C. Gell, V. Bormuth, and J. Howard. 2011b. Depolymerizing kinesins Kip3 and MCAK shape cellular microtubule architecture by differential control of catastrophe. Cell. 147:1092–1103. doi:10.1016/j.cell.2011.10.037.

Gardner, M.K., M. Zanic, and J. Howard. 2013. Microtubule catastrophe and rescue. Curr. Opin. Cell Biol. 25:1–9. doi:10.1016/j.ceb.2012.09.006.

Gell, C., V. Bormuth, G.J. Brouhard, D.N. Cohen, S. Diez, C.T. Friel, J. Helenius, B. Nitzsche, H. Petzold, J. Ribbe, E. Schaffer, J.H. Stear, A. Trushko, V. Varga, P.O. Widlund, M. Zanic, and J. Howard. 2010. Microtubule dynamics reconstituted in vitro and imaged by single-molecule fluorescence microscopy. 95. First edit. Elsevier. 221–245 pp.

Gudimchuk, N.B., E. V Ulyanov, E.O. Toole, C.L. Page, D.S. Vinogradov, G. Morgan, G. Li, J.K. Moore, E. Szczesna, A. Roll-mecak, F.I. Ataullakhanov, and J.R. Mcintosh. 2020. Mechanisms of microtubule dynamics and force generation examined with computational modeling and electron cryotomography. Nat. Commun. 1–15. doi:10.1038/s41467-020-17553-2.

Guesdon, A., F. Bazile, R.M. Buey, R. Mohan, S. Monier, R.R. García, M. Angevin, C. Heichette, R. Wieneke, R. Tampé, L. Duchesne, A. Akhmanova, M.O. Steinmetz, and D. Chrétien. 2016. EB1 interacts with outwardly curved and straight regions of the microtubule lattice. Nat. Cell Biol. 1. doi:10.1038/ncb3412.

Hunter, A.W., M. Caplow, D.L. Coy, W.O. Hancock, S. Diez, L. Wordeman, and J. Howard. 2003. The kinesin-related protein MCAK is a microtubule depolymerase that forms an ATP-hydrolyzing complex at microtubule ends. Mol. Cell. 11:445–457. doi:10.1016/S1097-2765(03)00049-2.

Hyman, A.A., D.N. Drechsel, D. Kellogg, S. Salser, K. Sawin, P. Steffen, L. Wordeman, and T.J. Mitchison. 1991. Preparation of Modified Tubulins. Methods Enzymol. 196:478–485.

Kerssemakers, J.W.J., E.L. Munteanu, L. Laan, T.L. Noetzel, M.E. Janson, and M. Dogterom. 2006. Assembly dynamics of microtubules at molecular resolution. Nature. 442:709–712. doi:10.1038/nature04928.

Kim, T., and L.M. Rice. 2019. Long-range, through-lattice coupling improves predictions of microtubule catastrophe. Mol. Biol. Cell. 30:mbc.E18-10-0641. doi:10.1091/mbc.E18-10-0641.

Lawrence, E.J., G. Arpag, S.R. Norris, and M. Zanic. 2018. Human CLASP2 specifically regulates microtubule catastrophe and rescue. Mol. Biol. Cell. 29:1168–1177. doi:10.1091/mbc.E18-01-0016.

Mandelkow, E.-M., E. Mandelkow, and R.A. Milliganll. 1991. Microtubule dynamics and microtubule caps: a time-resolved cryo-electron microscopy study. J. Cell Biol. 114:977–991. doi:10.1083/jcb.114.5.977.

Margolin, G., I. V. Gregoretti, T.M. Cickovski, C. Li, W. Shi, M.S. Alber, and H. V. Goodson. 2012. The mechanisms of microtubule catastrophe and rescue: Implications from analysis of a dimer-scale computational model. Mol. Biol. Cell. 23:642–656. doi:10.1091/mbc.E11-08-0688.

Maurer, S.P., F.J. Fourniol, G. Bohner, C.A. Moores, and T. Surrey. 2012. EBs recognize a nucleotide-dependent structural cap at growing microtubule ends. Cell. 149:371–382. doi:10.1016/j.cell.2012.02.049.

Mcintosh, J.R., E.O. Toole, G. Morgan, J. Austin, E. Ulyanov, F.I. Ataullakhanov, and N.B. Gudimchuk. 2018. Microtubules grow by the addition of bent guanosine triphosphate tubulin to the tips of curved protofilaments. J. Cell Biol. 1–25.

Michaels, T.C.T., S. Feng, H. Liang, and L. Mahadevan. 2020. Mechanics and kinetics of dynamic instability. Elife. e54077. doi:10.7554/eLife.54077.

Mickolajczyk, K.J., E.A. Geyer, T. Kim, L.M. Rice, and W.O. Hancock. 2019. Direct observation of individual tubulin dimers binding to growing microtubules. PNAS. 1–15. doi:10.1101/418053.

Mimori-Kiyosue, Y., I. Grigoriev, G. Lansbergen, H. Sasaki, C. Matsui, F. Severin, N. Galjart, F. Grosveld, I.A. Vorobjev, S. Tsukita, and A. Akhmanova. 2005. CLASP1 and CLASP2 bind to EB1 and regulate microtubule plus-end dynamics at the cell cortex. J. Cell Biol. 168:141–153. doi:10.1083/jcb.200405094.

Mitchison, T.J., and M.W. Kirschner. 1984. Dynamic instability of microtubule growth. Nature. 310:237–242.

Odde, D.J., L. Cassimeris, and H.M. Buettner. 1995. Kinetics of microtubule catastrophe assessed by probabilistic analysis. Biophys. J. 69:796–802. doi:10.1016/S0006-3495(95)79953-2.

Orbach, R., and J. Howard. 2019. The dynamic and structural properties of axonemal tubulins support the high length stability of cilia. Nat. Commun. 10:1–11. doi:10.1038/s41467-019-09779-6.

Prahl, L.S., B.T. Castle, M.K. Gardner, and D.J. Odde. 2014. Quantitative analysis of microtubule self-assembly kinetics and tip structure. 540. 1st ed. Elsevier Inc. 35–52 pp.

Reid, T.A., C. Coombes, S. Mukherjee, R.R. Goldblum, K. White, S. Parmar, M. Mcclellan, M. Zanic, N. Courtemanche, and M.K. Gardner. 2019. Structural state recognition facilitates tip tracking of EB1 at growing microtubule ends. Elife. 1–32. doi:10.1101/636092.

Rickman, J., C. Duellberg, N.I. Cade, L.D. Griffin, and T. Surrey. 2017. Steady-state EB cap size fluctuations are determined by stochastic microtubule growth and maturation. Proc. Natl. Acad. Sci. 201620274. doi:10.1073/pnas.1620274114.

Roostalu, J., C. Thomas, N.I. Cade, S. Kunzelmann, I.A. Taylor, and T. Surrey. 2020. The speed of GTP hydrolysis determines GTP cap size and controls microtubule stability. Elife. 9:1–22. doi:10.7554/eLife.51992.

Ruhnow, F., D. Zwicker, and S. Diez. 2011. Tracking single particles and elongated filaments with nanometer precision. Biophys. J. 100:2820–2828. doi:10.1016/j.bpj.2011.04.023.

Rusan, N.M., C.J. Fagerstrom, A.-M.C. Yvon, and P. Wadsworth. 2001. Cell cycle-dependent changes in microtubule dynamics in living cells expressing green fluorescent protein-alpha tubulin. Mol. Biol. Cell. 12:971–80.

Schek, H.T., M.K. Gardner, J. Cheng, D.J. Odde, and A.J. Hunt. 2007. Microtubule Assembly Dynamics at the Nanoscale. Curr. Biol. 17:1445–1455. doi:10.1016/j.cub.2007.07.011.

Schindelin, J., I. Arganda-Carreras, E. Frise, V. Kaynig, M. Longair, T. Pietzsch, S. Preibisch, C. Rueden, S. Saalfeld, B. Schmid, J.Y. Tinevez, D.J. White, V. Hartenstein, K. Eliceiri, P. Tomancak, and A. Cardona. 2012. Fiji: An open-source platform for biological-image analysis. Nat. Methods. 9:676–682. doi:10.1038/nmeth.2019.

Slep, K.C. 2009. The role of TOG domains in microtubule plus end dynamics. Biochem. Soc. Trans. 37:1002–1006. doi:10.1042/BST0371002.

Strothman, C., V. Farmer, G. Arpağ, N. Rodgers, M. Podolski, S. Norris, R. Ohi, and M. Zanic. 2019. Microtubule minus-end stability is dictated by the tubulin off-rate. J. Cell Biol. 218:2841–2853. doi:10.1083/jcb.201905019.

Tarantino, N., J.Y. Tinevez, E.F. Crowell, B. Boisson, R. Henriques, M. Mhlanga, F. Agou, A. Israel, and E. Laplantine. 2014. Tnf and il-1 exhibit distinct ubiquitin requirements for inducing NEMO-IKK supramolecular structures. J. Cell Biol. 204:231–245. doi:10.1083/jcb.201307172.

VanBuren, V., L. Cassimeris, and D.J. Odde. 2005. Mechanochemical model of microtubule structure and self-assembly kinetics. Biophys. J. 89:2911–2926. doi:10.1529/biophysj.105.060913.

VanBuren, V., D.J. Odde, and L. Cassimeris. 2002. Estimates of lateral and longitudinal bond energies within the microtubule lattice. Proc. Natl. Acad. Sci. 99:6035–6040. doi:10.1073/pnas.092504999.

Vasquez, R.J., D.L. Gard, and L. Cassimeris. 1994. XMAP from Xenopus Eggs Promotes Rapid Plus End Assembly of Microtubules and Rapid Microtubule Polymer Turnover. J. Cell Biol. 127:985–993.

Vitre, B., F.M. Coquelle, C. Heichette, C. Garnier, D. Chrétien, and I. Arnal. 2008. EB1 regulates microtubule dynamics and tubulin sheet closure in vitro. Nat. Cell Biol. 10:415–421. doi:10.1038/ncb1703.

Walker, R.A., E.T. O’Brien, N.K. Pryer, M.F. Soboeiro, W.A. Voter, H.P. Erickson, and E.D. Salmon. 1988. Dynamic Instability of Individual Microtubules. J. Cell Biol. 107:1437–1448.

Zakharov, P., N.B. Gudimchuk, V. Voevodin, A. Tikhonravov, F.I. Ataullakhanov, and E.L. Grishchuk. 2015. Molecular and Mechanical Causes of Microtubule Catastrophe and Aging. Biophys. J. 109:2574–2591. doi:10.1016/j.bpj.2015.10.048.

Zanic, M. 2016. Measuring the Effects of Microtubule-Associated Proteins on Microtubule Dynamics In Vitro. Methods Mol. Biol. 1413:47–61. doi:10.1007/978-1-4939-3542-0_4.

Zanic, M., J.H. Stear, A.A. Hyman, and J. Howard. 2009. EB1 recognizes the nucleotide state of tubulin in the microtubule lattice. PLoS One. 4:1–5. doi:10.1371/journal.pone.0007585.

Zanic, M., P.O. Widlund, A.A. Hyman, and J. Howard. 2013. Synergy between XMAP215 and EB1 increases microtubule growth rates to physiological levels. Nat. Cell Biol. 15:688–693. doi:10.1038/ncb2744.

Zhang, R., G.M. Alushin, A. Brown, and E. Nogales. 2015. Mechanistic origin of microtubule dynamic instability and its modulation by EB proteins. Cell. 162:849–859. doi:10.1016/j.cell.2015.07.012.

